# Deep grey matter volume loss drives disability worsening in multiple sclerosis

**DOI:** 10.1101/182006

**Authors:** Arman Eshaghi, Ferran Prados, Wallace Brownlee, Daniel R. Altmann, Carmen Tur, M. Jorge Cardoso, Floriana De Angelis, Steven H. van de Pavert, Niamh Cawley, Nicola De Stefano, M. Laura Stromillo, Marco Battaglini, Serena Ruggieri, Claudio Gasperini, Massimo Filippi, Maria A. Rocca, Alex Rovira, Jaume Sastre-Garriga, Hugo Vrenken, Cyra E Leurs, Joep Killestein, Lukas Pirpamer, Christian Enzinger, Sebastien Ourselin, Claudia A.M. Gandini Wheeler-Kingshott, Declan Chard, Alan J. Thompson, Daniel C. Alexander, Frederik Barkhof, Olga Ciccarelli (on behalf of the MAGNIMS study group)

**Author notes:** MAGNIMS steering committee members are listed in the appendix of this article.

## Abstract

**Objective:** Grey matter (GM) atrophy occurs in all multiple sclerosis (MS) phenotypes. We investigated whether there is a spatiotemporal pattern of GM atrophy that is associated with faster disability accumulation in MS.

**Methods:** We analysed 3,604 brain high-resolution T1-weighted MRI scans from 1,417 participants: 1,214 MS patients (253 clinically-isolated syndrome[CIS], 708 relapsing-remitting[RRMS], 128 secondary-progressive[SPMS], 125 primary-progressive[PPMS]), over an average follow-up of 2.41 years (standard deviation[SD]=1.97), and 203 healthy controls (HCs) [average follow-up=1.83 year, SD=1.77], attending 7 European centres. Disability was assessed with the Expanded-Disability Status Scale (EDSS). We obtained volumes of the deep GM (DGM), temporal, frontal, parietal, occipital and cerebellar GM, brainstem and cerebral white matter. Hierarchical mixed-models assessed annual percentage rate of regional tissue loss and identified regional volumes associated with time-to-EDSS progression.

**Results:** SPMS showed the lowest baseline volumes of cortical GM and DGM. Of all baseline regional volumes, only that of the DGM predicted time-to-EDSS progression (hazard ratio=0.73, 95% CIs 0.65, 0.82; *p*<0.001): for every standard deviation decrease in baseline DGM volume, the risk of presenting a shorter time to EDSS worsening during follow-up increased by 27%. Of all longitudinal measures, DGM showed the fastest annual rate of atrophy, which was faster in SPMS (-1.45%), PPMS (-1.66%), and RRMS (-1.34%) than CIS (-0.88%) and HCs (-0.94%)[*p*<0.01]. The rate of temporal GM atrophy in SPMS (-1.21%) was significantly faster than RRMS (-0.76%), CIS (-0.75%), and HCs (-0.51%). Similarly, the rate of parietal GM atrophy in SPMS (-1.24-%) was faster than CIS (-0.63%) and HCs (-0.23%) (all p values <0.05). Only the atrophy rate in DGM in patients was significantly associated with disability accumulation (beta=0.04, *p*<0.001).

**Interpretation:** This large multi-centre and longitudinal study shows that DGM volume loss drives disability accumulation in MS, and that temporal cortical GM shows accelerated atrophy in SPMS than RRMS. The difference in regional GM atrophy development between phenotypes needs to be taken into account when evaluating treatment effect of therapeutic interventions.

## Introduction

The clinical course of multiple sclerosis (MS) is heterogeneous. Some patients experience relapses with recovery (relapsing-remitting [RR] MS), while others develop progressive disability either from the onset (primary-progressive [PP] MS), or after a period of relapses (secondary-progressive [SP] MS). RRMS patients account for approximately 90% of cases at onset^1^, whose majority later progress to SPMS. The pathogenic mechanisms driving accrual of disability are beginning to be elucidated^2^: neurodegeneration plays a crucial role in determining accrual of disability over time^3^.

Neurodegeneration is reflected *in-vivo* by reduced brain volume (or brain atrophy), which can be measured by MRI^3^. Over time, brain volume declines more rapidly in MS patients when compared with age-matched healthy controls (HCs)^3–6^. Across MS phenotypes, SPMS shows the fastest annual rate of brain atrophy, which is estimated to be 0.6% (compared to about 0.2% in age-matched HCs)^5^. The role of brain atrophy in monitoring response to treatments in MS is evolving: whole brain atrophy has been recently used as primary outcome measure in Phase II clinical trials in SPMS^7,8^.

Whole brain atrophy is mainly driven by neuroaxonal loss in the GM^3^. GM volume loss is associated with long-term disability^9,10^, and explains physical disability better than white matter^9,11^ and whole brain atrophy^5^. Some GM regions, such as the cingulate cortex and thalamus, are affected by volume loss more extensively than others^12,13^, and the extent of their volume loss correlates with disability^13,14^, and cognitive impairment^15^. Regional predilection for atrophy is not unique to MS; for example, hippocampal atrophy is more pronounced than the whole brain atrophy in the early phase of Alzheimer’s disease^16^.Although cross-sectional studies have previously shown patterns of regional atrophy in different types of MS^12,17^, studies on longitudinal evolution of atrophy in different structures across MS phenotypes are scarce.

The overarching goal of our study was to investigate whether there is a spatiotemporal pattern of GM atrophy that is associated with faster disability accumulation in MS. In a large multi-centre cohort, which included all MS phenotypes and HCs, we tested the following hypotheses: (i) some GM regions show faster atrophy rate than others and their rate may differ between MS phenotypes; (ii) smaller baseline volumes of brain structures, reflecting a more extensive neurodegeneration, predict disability accrual; (iii) the rate of regional volume loss is associated with the rate of disability accumulation.

## Methods

### Participants

In this retrospective study we collected data from 7 European MS centres (MAGNIMS: www.magnims.eu) from 1,424 participants who have been studied between 1996 and 2016; we included participants who fulfilled the following criteria: (1) a diagnosis of MS according to 2010 McDonald Criteria^18^ or a clinically isolated syndrome (CIS)^19^; (2) healthy controls (HCs) without history of neurological or psychiatric disorders; (3) at least two-MRI scans acquired with a minimal interval of 6 months with identical protocol, including high-resolution T1-weighted MRI (allowing regional grey and white matter segmentation), and T2/Fluid Attenuated Inversion Recovery (FLAIR), sequences. Patients were scored on Expanded Disability Status Scale (EDSS)^20^. To increase the number of HCs scans, which were provided by 4 centres, we collected data from age-matched HCs from the Parkinson’s Progression Marker’s Initiative (http://www.ppmi-info.org/data).

MRI scans were taken under consent obtained from each subject independently in each centre. The final protocol for this study was reviewed and approved by the European MAGNIMS collaboration for analysis of pseudo-anonymised scans.

### Image acquisition

We included scans from 13 different MRI protocols; all centres except one provided 3D-T1 weighted scans (**Supplementary Table 1** and **Supplementary Table 2** show the MRI protocols).

### Image analysis

We performed image analysis as follows:

#### 1) Bias field correction

We used N4 bias field correction to correct for field inhomogeneity in T1-weighted scans using ANTs v2.10^21^.

#### 2) Lesion filling

Lesion masks were manually delineated on PD/T2 images by different raters at each centre semi-automatically, except for 3 centres that used the same automatic lesion segmentation with LST toolbox (version 2.0.15) ^22^.We calculated linear transformation matrices to register T2/FLAIR with the T1-weighted scan using FSL-FLIRT v5.0^23^. Then we applied these matrices to lesion masks to transfer them into the accompanying T1 subject-space. We used the FSL lesion filling method which uses a white matter mask calculated with FSL-FAST^24^ to fill T1 hypo-intensities within normal-appearing whiter matter, so to reduce segmentation errors, as previously done^25–27^.

#### 3) Symmetric within-subject registration

To avoid asymmetric registration and interpolation of longitudinal scans (e.g., toward the baseline scan), we constructed an unbiased subject-specific template that has “equal distance“ from each time point using FreeSurfer version 5.3^28–30^. We linearly transformed T1-weighted images to this symmetric space with the unbiased transformation matrix for each time point and used cubic B-spline interpolation to reduce interpolation artefacts. We manually checked the alignment of scans in the symmetric space.

#### 4) Tissue segmentation

Next, in the symmetric space, we segmented T1 scans into the GM, white matter and cerebrospinal fluid with the Geodesic Information Flow (GIF) software (part of NifySeg,http://cmictig.cs.ucl.ac.uk/niftyweb/program.php?p=GIF)^31^, and parcellated each hemisphere into regions of interest according to the Neuromorphometric atlas^32^. GIF uses an atlas propagation and label fusion strategy to calculate the voxel probabilities of GM, white matter and CSF^31^; this method has been previously used in MS and other neurodegenerative disorders^33,34^. The template library had 95 MRI brain scans (HCs and patients with Alzheimer’s disease) with neuroanatomic labels (http://www.neuromorphometrics.com/). This atlas, which is similar to Mindboggle atlas, was developed to improve the consistency and clarity of Desikan-Killiany protocol^32^.

To calculate brain masks and exclude segmentation errors outside of the brain we used STEPS (Similarity and Truth Estimation for Propagated Segmentations, http://cmictig.cs.ucl.ac.uk/niftyweb/program.php?p=BRAIN-STEPS) based on a template library of 682 hand-drawn brain masks^35,36^. These maps were applied to each time point separately.

#### 5) Regional volume calculation

We visually assessed the segmentations to assure the quality for statistical analysis. To calculate regional volumes, we summed the probability of the segmented tissue voxels (GM or white matter) in each parcellated region and multiplied the sum with the voxel volume. We averaged values between left and right hemispheres. Next, we summarised the regional volumes according to Neuromorphometrics protocol by summing the volume of GM regions in the temporal, parietal, occipital, frontal lobes, cerebellum and deep GM (DGM) [thalamus, putamen, globus pallidus, caudate, and amygdala]. We also obtained the volume of the brainstem and of the cerebral white matter.

**Figure 1** shows the image analysis pipeline.

**Figure 1.**
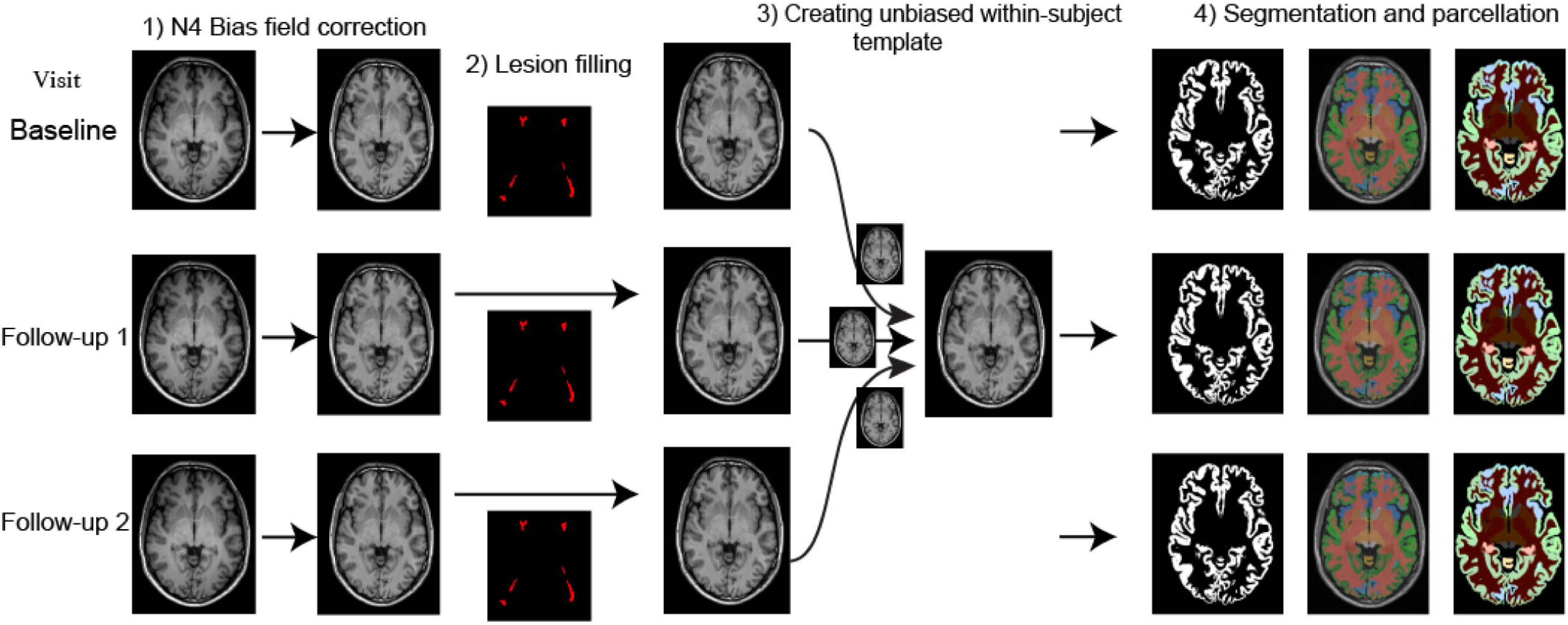
Imaging analysis pipeline. An unbiased symmetric image registration approach was used to calculate atrophy.

### Statistical analysis

#### Brain volumes at baseline and rates of volume changes over time

To investigate baseline volumes (intercept) and rates (slopes) of volume change by subject group and region, we used linear mixed-effects models with the volume at a given time as the response variable, and time and interactions with time as fixed-effect covariates^37^. This model estimates adjusted rate while allowing for nested correlation structures, such as time of visit within subject within scanner, by incorporating, in this example, subject and scanner random intercepts, and a random slope on time. The interaction terms with time (e.g., subject group X time), allows the estimation of rate differences across the interacting variable, in this example subject groups or clinical phenotypes. Including another interaction with time, such as gender X time, adjusts the rate for gender. In addition to time, the fixed-effect covariates were: scanner magnetic field, subject group, gender, age at baseline, total intracranial volume (sum of the volumes of GM, WM and CSF) at baseline; and the interactions of each of these with time. Disease duration was too highly correlated with age at baseline to give reliable estimation, and was omitted from the final models. To estimate the percentage changes per unit (year) increase in time, we log-transformed the volume^38^. We adjusted time to zero for those visits in which a patient converted from one phenotype to another (e.g., CIS to RRMS). We performed *post-hoc* analyses to identify specific GM regions within the cerebral lobes and among the DGM nuclei that showed significant differences between MS phenotypes, as well as the default-mode network regions^39^.

To investigate whether there is an association between the rate of loss in specific regions and MS phenotypes, 3-way interactions were used, for example, clinical phenotype × region × time. We used R (version 3.2.2) and the NLME package^40,41^.

For each model, we visually checked the heteroscedasticity (which is the unequal variance of a variable across the range of values of a second variable that predicts it) per group by plotting residuals against the fitted values.

We corrected for multiple comparisons accounting for the number of all the tests performed with the false-discovery rate method.

#### Effect of MRI protocols on imaging measures

To assess the effect of the MRI protocol on MRI measures (we took into account the protocols rather than the centres because some centres acquired more than one protocol with more than one scanner) we included it as a fixed-effect variable in a separate mixed-effect model, and calculated the average effect sizes for MRI protocols and MS phenotypes (i.e., disease effects) while fixing other variables.

#### Assessing associations between brain tissue volumes and disability accrual

For easier interpretation of clinical and imaging measures, we standardised volumes by subtracting the mean and dividing by the standard deviation (*Z*-score). We analysed CIS and relapse-onset patients together, because some patients had converted from CIS to RRMS, or from RRMS to SPMS. This allowed us to take advantage of a longer follow-up period. With similar mixed-effects models we investigated the following three questions: (1) Are the baseline volumes of the DGM, the temporal, frontal, parietal, occipital and cerebellar GM, brainstem, and white matter, and white matter lesion load associated with EDSS at baseline? (2) Are changes in all these regional volumes and white matter lesion load associated with EDSS changes over time? (3) Do baseline volumes of all these regions and white matter lesion at baseline predict time-to-EDSS progression (event=EDSS progression) during follow-up? The EDSS-progression event was defined as 1.5 increase in EDSS, if the baseline EDSS was 0; one-point increase if EDSS was less than or equal to 6; and 0.5 increase if EDSS was more than 6^42^. We used a Cox-regression model to explore whether baseline volumes of these structures predicted time to event. We performed a *post-hoc* analysis using all GM regions to determine the most important predictors of time-to-EDSS-progression (as defined above) and confirm that the results of the DGM were not affected by the bias of merging a higher number of cortical regions into the main lobes. We performed FDR correction to adjust for multiple comparisons.

#### Additional analyses: software reliability and effects of disease modifying treatments

We carried out additional analyses to assess the reliability of brain volumes estimated with GIF software, FSL-FIRST, and SPM12, and effects of treatments on atrophy measures (see **Supplemental Material**).

## Results

The MRI scans of 1,417 subjects were analysed (scans of three subjects were excluded due to significant motion artefacts on visual inspection and four due to registration issues because of missing MRI header information); 1,214 patients (253 had CIS, 708 had RRMS, 128 had SPMS, and 125 had PPMS), and 203 were HCs. In total, we analysed 3,604 T1-weighted MRI. Average number of scans per subject was 2.54 (SD=1.04), with an average follow up of 2.41years (SD=1.97) for patients, and 1.83 (SD=1.77) years for HCs (see **Table 1** for follow-up information per group). The total numbers of participants with 3 or more visits for each group were: 90 HCs, 48 CIS, 334 RRMS, 39 SPMS, and 58 PPMS. A total of 96 patients with CIS (38%) converted to RRMS, and 28 patients with RRMS (4%) converted to SPMS during the follow-up.

**Table 1.** Baseline characteristics of participants Group Healthy controls

| Group | Healthy<br>controls | CIS | RRMS | SPMS | PPMS |
| --- | --- | --- | --- | --- | --- |
| **Total number (number**<br>**of females)** | 203 (112) | 253 (171) | 708 (473) | 128 (75) | 125 (55) |
| **Average follow up in**<br>**years (range)** | 1.83 (0.5-7.8) | 1.46 (0.5-13) | 2.72 (0-13) | 2.06 (0-5.5) | 2.85 (0.5-6) |
| **Average age (± SD)** | 38.7 ± 10.5 | 33 ± 8 | 38.2 ± 9.8 | 48.2 ± 9.8 | 48.5 ± 10.1 |
| **Average disease**<br>**duration (± SD)** | — | 0.4 ± 1.4 | 6.7 ± 7.3 | 15.6 ± 9.9 | 6.8 ± 5.9 |
| **Median EDSS (range)** | — | 1 (0-4.5) | 2 (0-7) | 6 (2.5-9) | 5 (2-8) |
| **Median T2 lesion load**<br>**(ml) (1<sup>st</sup>-3<sup>rd</sup> quartiles)** | — | 2.97<br>(1.01-5.04) | 5.05<br>(2.05-11.79) | 11.04<br>(3.18-23.14) | 9.38<br>(2.69-22.02) |
| **% (number) of patients**<br>**on DMTs** | — | 20%<br>(52) | 49%<br>(345) | 41%<br>(52) | 6%<br>(8) |

There was a significant difference in gender ratio between groups (*p*<0.001, see Table 1 for gender ratios). Patients with progressive MS (SPMS and PPMS) had significantly greater disability than patients with RRMS and CIS (Mann-Whitney tests, *p*<0.001, see **Table 1**), and were older than RRMS (p<0.001, average difference=10.7 years), CIS (*p*<0.01, average difference=15.6 years) and HCs (*p*<0.01, average difference=10 years). Age was similar between patients with RRMS and HCs. Patients with CIS were younger than HCs (*p*<0.01, average difference=4.9 years). Patients with CIS had the lowest T2 lesion load, and patients with SPMS had the highest T2 lesion load. About half of patients with RRMS were on disease modifying treatments (see **Table 1**).

### Brain atrophy at baseline in MS and rates of volume changes over time

At baseline, all clinical phenotypes (CIS, RRMS, SPMS, and PPMS) had significantly smaller cortical GM and DGM volumes than HCs. SPMS showed the lowest cortical GM and DGM volumes, followed by PPMS, RRMS, CIS. All clinical phenotypes, but not CIS, had significantly reduced whole brain and white matter volumes when compared to HCs (see **Figure 2A**).

**Figure 2.**
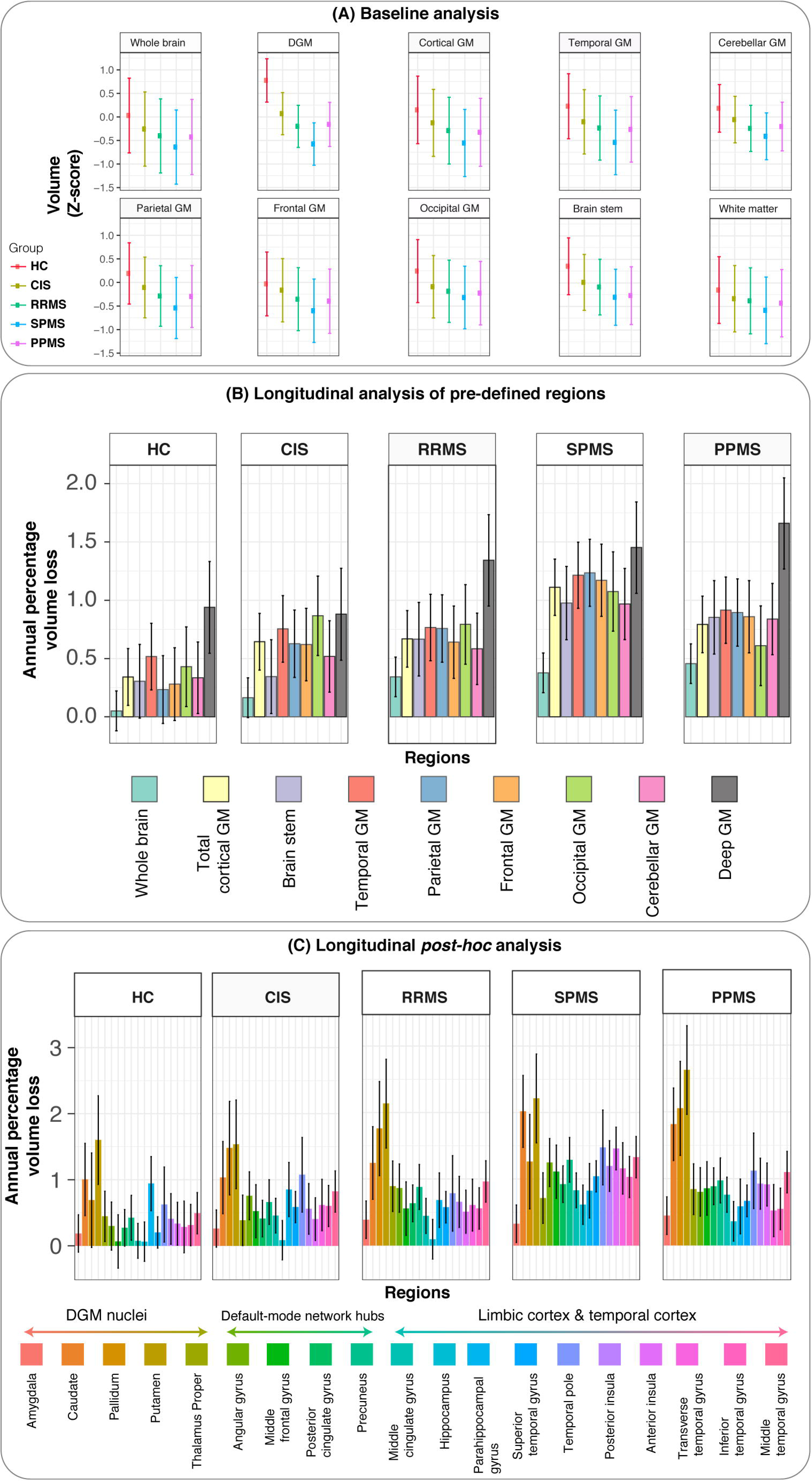
Baseline volumes, and annual percentage loss of brain regions in clinical phenotypes and healthy controls. Adjusted baseline values for HCs, CIS, RRMS, SPMS, and PPMS are shown in (A), where the adjusted mean is shown as a point, and error bars show the 95% confidence-interval. Adjusted *P*-values of pairwise comparisons between groups are shown in Supplementary Table 4. Longitudinal analyses are shown in (B) and (C). Bar charts of the adjusted annual percentage of loss are shown in (B) for the predefined regions. Height of each bar chart is the average estimate of the percentage annual loss from the mixed-effects model for each group. The error bars represent 95% confidence interval of these estimates. Adjusted *P*-values for pairwise comparison between regions across clinical phenotypes and HCs are shown in Supplementary Table 4. White matter volumes are not shown in (B, and C) because they did not show a significant change over time in any clinical phenotype. *Post-hoc* analyses of annual percentage loss are shown in (C) where DGM nuclei, temporal, limbic and default mode network regions were selected. Similar to (B) the adjusted average annual percentage volume loss for these regions is the height of each bar-chart and error bars represent 95% confidence intervals. Baseline values (A) and rates (B, and C) were adjusted in a single mixed-effects hierarchical model including age, gender, total intracranial volume at baseline, scanner magnetic field, and their interactions with time as the fixed-effects. Centre, subject and visits were nested (hierarchical) random-effects. Abbreviations: HC, healthy controls; CIS, clinically isolated syndrome; RRMS, relapsing-remitting multiple sclerosis; SPMS, secondary-progressive multiple sclerosis; PPMS, primary-progressive multiple sclerosis.

The fastest regional decline in tissue volume over time was seen in the DGM in all clinical phenotypes (PPMS: −1.66% per year, SPMS: −1.45%, RRMS: −1.34%, CIS: −0.88%, *p*<0.01) and in HCs (-0.94%). The rate of atrophy in the DGM was greater in RRMS, SPMS and PPMS than CIS and HCs (all p values <0.01) (**Figure 2B** and **Supplementary Tables 3 and 4**), but did not differ between RRMS, SPMS and PPMS. The rate of volume loss in the DGM in all MS patients together was significantly higher than that in the cortical and cerebellar GM and brainstem (although the rate of volume loss over time in these areas was still significant) (all p values < 0.05).

The volume loss of the whole cortical GM was faster in SPMS (-1.11% per year), PPMS (-0.79%), RRMS (-0.67%), than HCs (-0.34%)(all p values <0.05). Among the cortical regions, the temporal lobe GM showed a faster volume loss in SPMS (-1.21%) than RRMS (-0.77%) and CIS (-0.75%) (all p values <0.05) (**Figure 2B and Supplementary Tables 3 and 4**). Similarly, the parietal GM showed a faster volume loss in SPMS (-1.24%) than CIS (-0.63%) (p<0.05) (**Figure 2B and Supplementary Tables 3 and 4**). No differences in rates of volume loss were seen in the frontal and occipital GM between clinical phenotypes. Overall, all the cortical GM regions, with the exception of the occipital cortex, showed a faster rate of atrophy in MS than HCs (**Figure 2B** and **Supplementary Table 4**).

The white matter did not show a significant rate of volume loss in HCs or any of the clinical phenotypes.

There was no heteroscedasticity in the plots of residuals against fitted values.

In the *post-hoc* analyses when looking at regions and clinical phenotypes we found that among the DGM nuclei, the putamen showed the fastest volume loss in PPMS (-2.6%).Within the temporal lobe GM, the fastest volume loss was seen in the temporal pole (-1.47%) and posterior insula in SPMS (-1.19%). When looking at the parietal lobe GM, the precuneus showed the fastest atrophy rates in SPMS (-1.28%) (**Figure 2C**). Whilst the fastest rate of atrophy was seen in DGM in SPMS, the temporal lobe GM showed the highest difference between SPMS and HCs (see **Figure 2C**).

There was no significant effect of gender on rates of atrophy. There was no significant association between GM volumes and T2 (or FLAIR) lesion load.

### Regions showing the highest rate of loss

When we compared the rate of volume loss across different regions in all patients (CIS, RRMS, SPMS, and PPMS) together, the fastest decline (or lowest slope) was seen in the DGM (**Supplementary Tables 3** and **4**). The rate of loss in the cortical GM regions was similar between lobes and to that of the cerebellum. The slowest rate of loss was seen in the brainstem.

### Spatiotemporal pattern of GM volume loss in clinical phenotypes

Although SPMS showed the lowest baseline volumes of cortical GM and DGM, and the rate of the DGM volume loss was faster in SPMS, PPMS and RRMS than CIS and HCs, there was no significant association between the rate of loss in specific regions and clinical phenotypes, which suggests that all clinical phenotypes share a similar spatiotemporal pattern of GM loss.

### Effect of MRI protocols on imaging measures

The average effects of MS phenotypes on brain volumes at baseline were higher than the protocol effect on the brain volumes (protocol effects: whole brain = 4.3%, cortical GM =5.1%, DGM = 8.5%, disease effects: whole brain = 4.8%, cortical GM =5.2%, DGM=13.7%). The average effects of MS phenotypes were higher than the effects of protocol on the rates of atrophy of the cortical GM and DGM (protocol effects: cortical GM = 0.14%, DGM = 0.21%, disease effects: cortical GM = 0.57%, DGM =0.53%), but not those of the whole brain (protocol effect = 0.51%, and disease effect = 0.38%).

### Association between EDSS and GM loss

In all clinical phenotypes combined, lower DGM and cortical GM volumes at baseline were associated with higher disability, as measured by the EDSS (*β*: DGM =-0.71, *p*<0.0001; cortical GM (*β*=-0.22, *p*<0.0001). Under the assumption of a linear relationship between EDSS and GM volume, this suggests that for every *Z*-score decrease in the DGM and cortical volume at baseline, the baseline EDSS increased on average by 0.7 and 0.22, respectively. There was a significant progression of EDSS in both relapse-onset and PPMS patients, which on average increased by 0.07 and 0.2 per year, respectively. When we examined associations between the rate of EDSS changes and rate of changes in the volumes of cortical GM regions, cerebellar GM and DGM over time, only the rate of loss in the DGM was associated with disability accumulation (*β*=-0.04, 95% CI: −0.02, −0.06, *p*=0.006). Under the assumption of a linear relationship between EDSS and rate of GM volume loss over time, this suggests that every standard deviation (*Z*-score) loss in the rate of DGM volume corresponded to an annual EDSS gain of 0.04.

The percentage of patients who had EDSS progression during follow-up (or who experienced the “event“) was 26%. When we looked at baseline predictors of disability accumulation, without any longitudinal imaging measure in the model, only the DGM predicted future EDSS progression. The hazard ratio [95% CI, *p*-value] for time-to-EDSS progression was 0.73 [95% CI 0.65, 0.82, *p*<0.0001], which suggests that for every standard deviation (*Z-*score) decrease in the DGM volume at baseline the risk of presenting a shorter time to EDSS worsening during the follow-up increased by 27% [95% CI: 18-35%]. The hazard ratio remained similar when we analysed relapse-onset and PPMS patients separately (0.72 and 0.73 respectively). **Figure 3** illustrates the survival-curve for these analyses.

**Figure 3.**
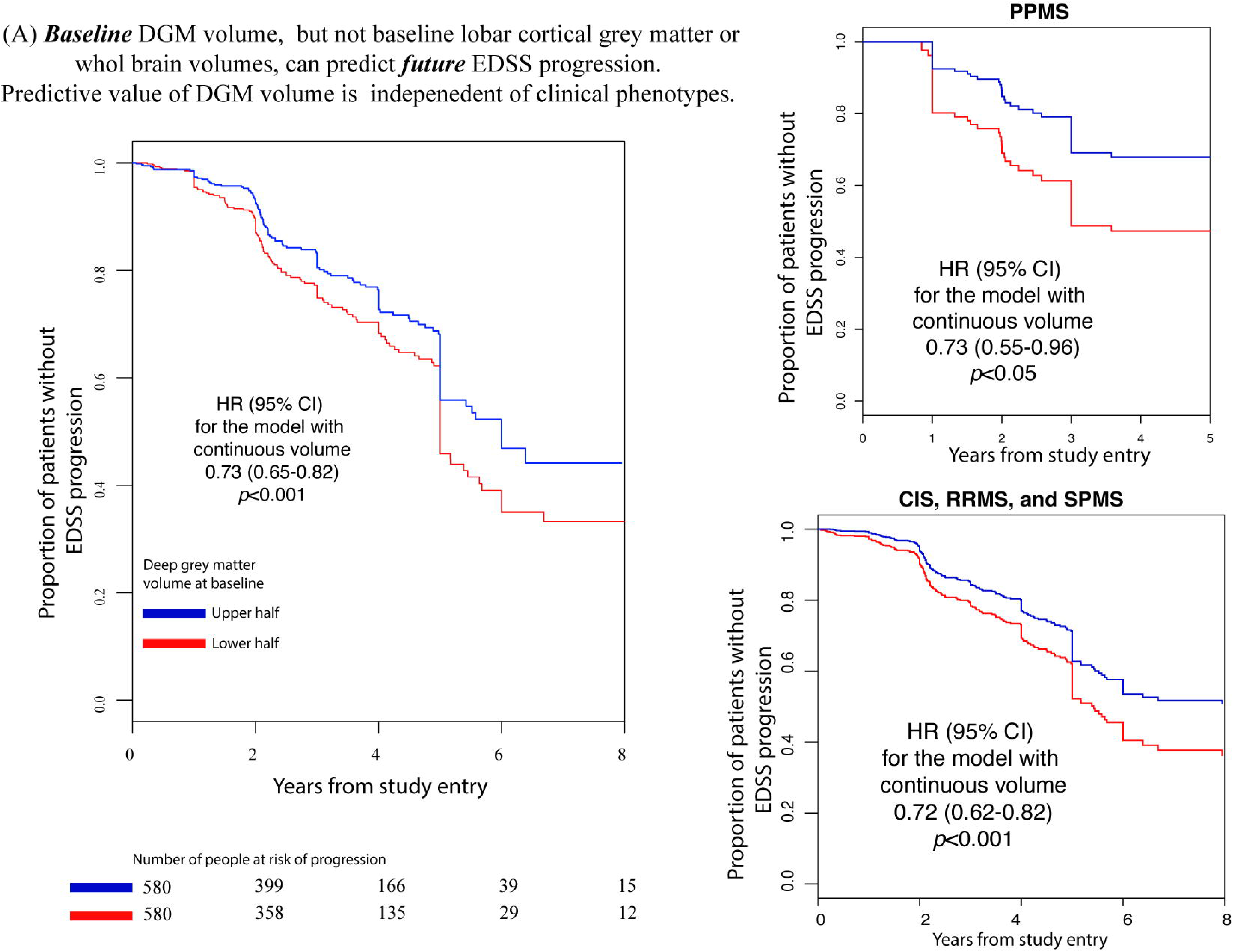
DGM volume predicts future progression of EDSS. Survival curves for time to event (sustained EDSS progression, see methods for definition) in CIS, relapse-onset and PPMS. We have analysed CIS and relapse-onset patient together, because a proportion of patients convert from CIS to RRMS, or from RRMS to SPMS during the course of study. Hazard-ratios for models with continuous outcome variables (regional volumes) are reported.

In the *post-hoc* analyses, baseline thalamic volume had the highest predictive value of EDSS-progression during follow-up in both PPMS and the relapse-onset groups, by increasing the risk to a shorter time to EDSS worsening of 37% in relapse-onset MS and 40% in PPMS (**Figure 4B** and **C**). In this analysis, the predictive value of the thalamus was followed by that of the hippocampus and angular gyrus in relapse-onset MS (**Figure 4B**), and by that of the putamen, posterior insula and temporal pole in PPMS (**Figure 4C**).

**Figure 4.**
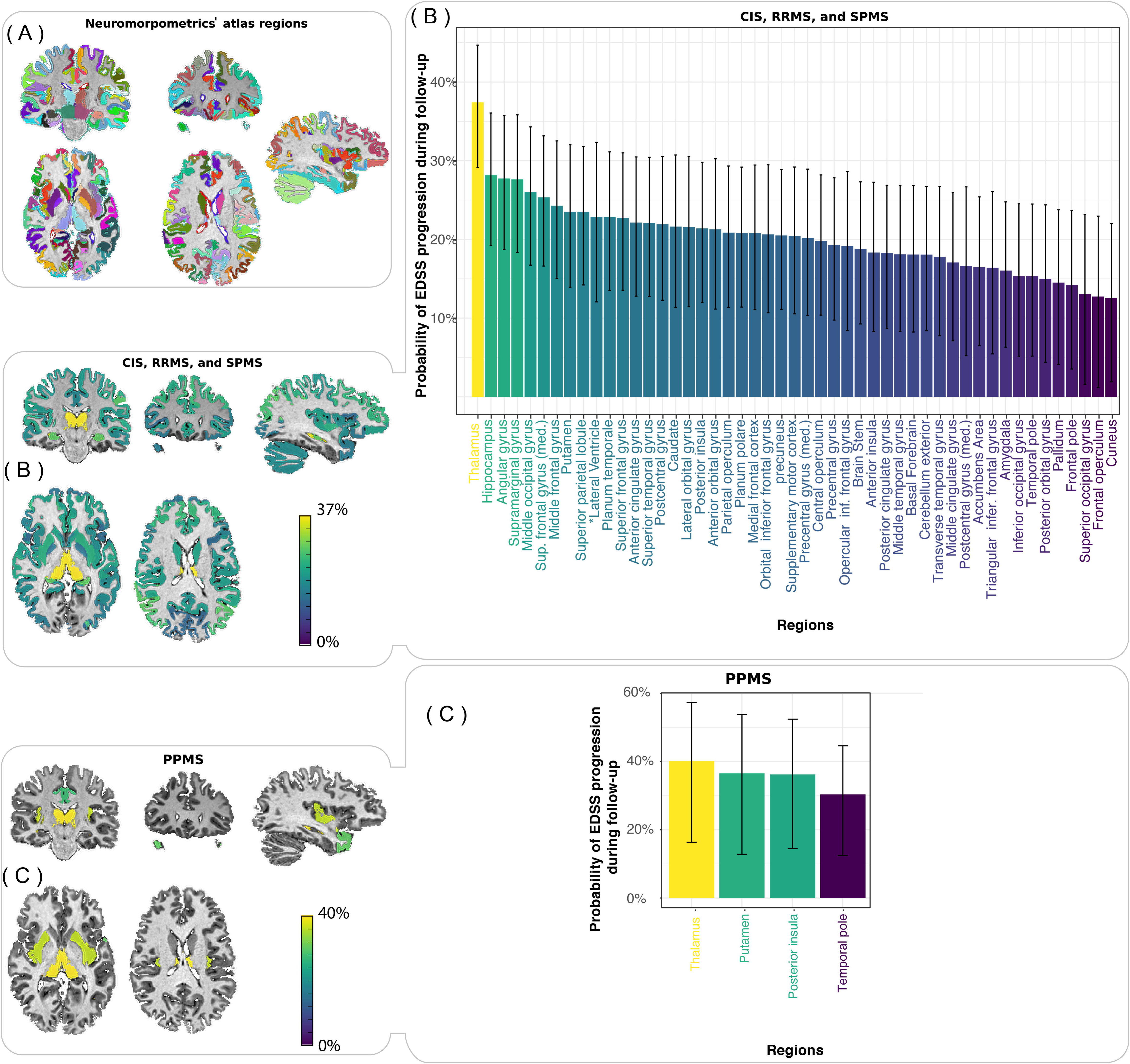
Risk of EDSS-progression during follow-up for each *Z*-score volume loss of the brain regions at baseline (*post-hoc* analysis). Results of the *post-hoc* Cox-Proportional Hazards univariate models are shown for the time-to-event analyses (event = sustained EDSS-worsening, see methods for the definition) in the regions of Neuromorphometrics’ atlas, which are shown in (A). The predictors were the baseline volumes of the regions shown in the *x*-axes of (B) for CIS, RRMS, and SPMS and (C) for PPMS. CIS, RRMS, and SPMS were analysed together, because several patients convert from one phenotype to another. Brain maps are shown in the left column, and bar-charts of the same analyses are shown in the right column of (B) and (C). Only regions whose *P*-value of the survival analysis survived FDR-correction (adjusted *P*<0.05) are shown in (B) and (C). The *y-*axes show the risk of progression for each *Z*-score loss in the volume of the corresponding brain region on *x*-axes. For example, for every *Z-*score loss of the thalamus volume at baseline, the risk of EDSS worsening during follow-up increased by 37% for the CIS, RRMS, SPMS group, and 40% for PPMS. Colour maps code the importance of baseline volumes of the regions to predict EDSS-worsening (or EDSS-progression) during follow-up. The absolute values of coefficients for ventricular volumes are shown in (B), because they have an effect in the opposite direction of other structures. Error-bars indicate the 95% confidence intervals.

There were no significant differences in the rates of loss in patients who were receiving disease-modifying drugs and those who were not (see **Supplementary Text**). The analyses with GIF software, FSL-FIRST, and SPM12 confirmed the reliability of brain volumes estimates (see details in **Supplementary Text**).

## Discussion

In this large multicentre study, we have shown that volume loss in DGM over time was faster than that seen in other brain regions across all clinical phenotypes, and DGM volume loss was the only GM region associated with disability accumulation. Additionally, we found that the smaller DGM volume at baseline was associated with increased risk of shorter time to EDSS progression, in agreement with previous studies that showed smaller DGM volume associated with higher disability^14,15^. Interestingly, we found that atrophy rates of the GM of cortical lobes were the fastest in SPMS, and were faster in the temporal lobe in SPMS in comparison with RRMS and CIS and in the parietal lobe in SPMS in comparison with CIS. However, no significant association between cortical regions and disability progression was detected. Overall, our findings suggest that the development of DGM atrophy may drive disability accumulation irrespective of clinical phenotypes, thereby becoming a useful outcome measure in neuroprotective clinical trials. Although the spatiotemporal pattern of atrophy remains similar across MS phenotypes, some cortical regions show accelerated atrophy in SPMS than RRMS and/or CIS. We now discuss these results in turn and in detail.

The pathological events that underpin DGM atrophy are not known, but this is generally interpreted as the result of neurodegeneration. Previous studies have shown that DGM atrophy is more severe in patients with progressive MS, longer disease duration and worse cognitive performance ^14,15,45^. Our *post-hoc* analyses showed that the thalamus, which is the DGM’s largest component, was a better predictor of future disability than other regions, and the rate of atrophy in the putamen was the highest across DGM nuclei. Previous studies, including those using advanced MRI, have found that thalamic damage at study entry was associated with higher disability^13–15^. DGM structures are extensively connected with cortical GM regions, and therefore DGM atrophy could be due to retrograde and anterograde neurodegeneration via tracts that connect GM areas. For example, the extent of cellular density loss in the thalamus, is associated with neurodegeneration in the remote (but connected) cortical regions, over and beyond the extent of atrophy explained by demyelination in connecting tracts^46^. There is also evidence of other neurodegenerative mechanisms in the DGM nuclei. For example, their higher load of iron than other regions can accumulate oxidised lipids which are associated with neurodegeneration^47^. In our healthy controls, the rate of DGM atrophy was faster than that in other regions, suggesting that it may be a hot spot for both age- and disease-related atrophy in the human brain, although a methodological issue, related to its more uniform structure than other brain regions, cannot be excluded. Nevertheless, the DGM volume holds strong promise as a marker of disease progression with the potential to respond to neuroprotective treatments that target neurodegeneration in MS.

Interestingly, the temporal lobe showed a significant acceleration in SPMS when compared to both RRMS and CIS. Similarly, the parietal lobe GM showed a significant acceleration of atrophy in SPMS in comparison with CIS. Our *post-hoc* analysis showed that the temporal pole and insula were the most affected structures in the temporal GM. Pathological studies have demonstrated an increase in the rate of neurodegeneration, especially in the temporal regions, during progressive stages of MS in comparison with RRMS and CIS^49,50^. Overall, a global pathological process in MS^51^, may become more pronounced in certain regions, such as the temporal GM, because of other mechanisms, such as static exposure to CSF (the insula in the temporal lobe) or hypoxia in watershed areas (some DGM nuclei such as the pallidum). For example, meningeal inflammation and cortical demyelination, which may play a role in cortical atrophy, preferentially affect deep sulci, such as the insula, where there is more exposure to static inflammatory cytokines^2,49,52^. Our findings also suggest that regions with more connections may be vulnerable to atrophy. For example, among the parietal cortical regions, the precuneus, a core part of an important functional brain network (default mode network), showed the fastest atrophy rates in SPMS^39^. Thus, acceleration of atrophy during SPMS may be explained by cortical network collapse with advancing of degeneration from initial injury sites (focal lesions in the white matter or initial DGM degeneration) to interconnected neocortical systems^53^. We found that MS phenotypes shared a common spatiotemporal pattern of volume loss (no significant 3-way interaction of time × region × phenotype). This shows, in line with previous studies, that the difference in pathology of progressive MS is only quantitative rather than qualitative in comparison with RRMS^2,55^.

Cortical GM atrophy was seen at study entry across clinical phenotypes, even in CIS, when compared with HCs, and was the greatest in progressive MS, in agreement with earlier studies^17,56^.Our findings of faster whole brain atrophy in SPMS, PPMS, RRMS than CIS, who in turn, showed higher cortical atrophy than HCs, are similar to previous studies on longitudinal whole brain atrophy^5,57,58^, regional atrophy^17,59–61^, and pathology of MS phenotypes^2,49^. Our study confirms our previous findings that relationships between whole brain atrophy and clinical changes are weak or absent^5^, and shows DGM atrophy as a stronger marker of clinical disability. Although the GM volumes of cortical lobes could not predict future EDSS progression, the more detailed *post-hoc* analyses showed that regional volumes, such those of the hippocampus and the angular gyrus, were associated with future EDSS progression. These regions are highly connected to other regions, and especially the angular gyrus (like the precuneus) acts as a hub in the default mode network, which could make it vulnerable to atrophy, as explained above^39^.

This study was not designed to assess the effect of treatment on atrophy rates, but does study atrophy while adjusting for possible confounding effects. The rates of atrophy in all clinical phenotypes were similar in people who were receiving disease-modifying treatments to those who were not. Even though we could not ascertain the duration of treatments due to retrospective nature of this study, the majority (90%) of patients on disease modifying treatments, were receiving first-line injectable drugs (interferon or glatiramer acetate) before study entry. The effects of these drugs on brain atrophy are modest at best ^62,63^. Therefore, drug effects are unlikely to be confounders of our analysis.

One strength of our study is that we included a large number of patients, who underwent the same protocol on the same MRI scanner over time at single sites. However, different MRI protocols could have an effect on atrophy measures and is a limitation of our study^64,65^. We therefore used a hierarchical statistical design based on scanner. Our study was powerful enough because the effects of clinical phenotype on the regional rates of atrophy were higher than the effects of between-centre variation.

We chose GIF software to segment and parcellate the brain^31^ because it allowed inclusion of 2D MRI data (which we had for one centre), and did not require any manual editing, unlike Freesurfer, which would have been unfeasible for such large number of scans. Our reliability analysis showed excellent agreement between GIF-derived DGM volume and that obtained using FSL-FIRST, and between GIF-derived cortical volumes and those obtained using SPM12, respectively. Therefore, we chose to present the results obtained with GIF because it allowed us to rely on only one method to segment DGM and cortical GM, and estimate TIV. We used TIV to adjust for variations in head size, rather than the skull-size, so that a more reliable estimate of head size is obtained, irrespectively of the field-of-view, the choice of the inferior cut-off of the brain for the analysis, and demographic factors (e.g., age, or weight)^70^. With regard to the statistical methods, we used mixed-effects models to calculate atrophy rates^41^, which naturally accommodated multiple (3 or more) time-points with varying intervals between follow-ups, and patients who convert from one phenotype to another (e.g., CIS to RRMS). These two issues are cumbersome to address with methods that rely on pairwise comparisons (e.g., SIENA, BSI) and suffer from higher variance in brain atrophy estimates as the interval between two scans increases^66,67^. Mixed-effects modelling, instead, estimates a variance component to eliminate implausible inconsistencies^68,69^. Based on our experience and the results of this study, we recommend the acquisition of high-resolution 3DT1 images (isotropic 1mm^3^). Several methods can calculate DGM volumes, such as FSL-FIRST, and GIF. We recommend the use of the GIF software when it is desirable to use the same method to segment both the cortex and DGM.

There were also limitations in this study. The majority of centres did not provide MRI scans of HCs, however, we included a large number of HCs including those from an external initiative (PPMI). Our findings of volume changes in HCs were consistent with the literature. Meta-analyses have shown, in individuals less than 70 years of age, rate of whole brain loss ranges from 0 to −0.5 (our study = −0.04), GM loss ranges from 0 to −0.5% per year (cortical GM in our study = −0.34%)^71^, and the subcortical structures may show loss of up to −1.12% (DGM in our study = −0.94)^72^. Cognitive functions were not tested, and it is unknown whether cortical patterns of GM atrophy over time were associated with cognitive impairment. Clinical trials in MS (and in progressive MS in particular) include confirmed disability progression, based on the EDSS, as primary outcome measure. Although for EDSS the model-estimated coefficients and their *p*-values and confidence intervals are valid for comparison between brain regions, the absolute value of these coefficients must be interpreted with caution, because the EDSS does not have a uniform linear interpretation. Since this was a retrospective study, the duration of treatments before entry to the study could not be ascertained for all participants. Disease modifying drugs may have lasting effects, for example they may slow the accrual of disability after a decade^63,73^. Moreover, MRI sequences sensitive to cortical lesions were not available, and the effects of cortical lesions on atrophy measures remain unknown.

In conclusion, the DGM atrophy showed the most rapid development over time– extending previous cross-sectional studies that showed a relationship between DGM atrophy and disability– was most closely associated with disability accumulation and predicted the time to EDSS worsening. In phase II trials of neuroprotective medications in MS, DGM atrophy measures may therefore have greater potential to show treatment effects than other regional GM or whole brain measures. There was a disconnect between DGM atrophy and cortical atrophy rates. The temporal and parietal cortices showed a faster rate of atrophy in SPMS than RRMS and/or CIS, whilst DGM showed a faster rater of atrophy in SPMS than CIS only, suggesting that neurodegeneration in GM regions may proceed at a different rate which should be taken into account in the design of clinical trials.

## Acknowledgements

A Eshaghi has received an ECTRIMS-MAGNIMS Fellowship (http://www.magnims.eu) and MSIF McDonald Fellowship (http://www.msif.org). Authors acknowledge The National Institute for Health Research (NIHR) Biomedical Research Centre (BRC) at University College London Hospitals (UCLH) for their support. C. Tur has received an ECTRIMS postdoctoral research fellowship in 2015. D. Alexander has received funding for this work from EPSRC (M020533, M006093, J020990) as well as the *European Union’s Horizon 2020 research and innovation programme* under grant agreement Nos 634541 and 666992. PPMI (http://www.ppmi-info.org) – a public-private partnership –is funded by the Michael J. Fox Foundation for Parkinson’s Research and funding partners (see http://www.ppmiinfo.org/about-ppmi/who-we-are/study-sponsors/ for the full list). This manuscript is under peer-review. We will provide the address to the journal website once it is published.

## Author Contributions

AE, DCA, OC, AJT, FB, FP, MJC, DC, DRA contributed to the conception and design of the study. OC, WB, CT, FDA, SHP, NC, NDS, MLS, MB, SR, CG, MF, MAR, AR, JSG, HV, CEL, JK, LP, and CE contributed to data acquisition and analysis. All authors contributed to drafting the text or preparing the figures.

## Potential Conflicts of Interest

J. Killestein has accepted speaker and consulting fees from Merck-Serono, Biogen, TEVA, Genzyme, Roche and Novartis.

N De Stefano has received honoraria from Biogen-Idec, Genzyme, Merck Serono, Novartis, Roche and Teva for consulting services, speaking and travel support. He serves on advisory boards for Merck Serono, Novartis, Biogen-Idec, Roche, and Genzyme, he has received research grant support from the Italian MS Society.

C Enzinger received funding for traveling and speaker honoraria from Biogen Idec, Bayer Schering Pharma, Merck Serono, Novartis, Genzyme and Teva Pharmaceutical Industries Ltd./sanofi-aventis; received research support from Merck Serono, Biogen Idec, and Teva Pharmaceutical Industries Ltd./sanofi-aventis; and serves on scientific advisory boards for Bayer Schering Pharma, Biogen Idec, Merck Serono, Novartis, Genzyme, Roche, and Teva Pharmaceutical Industries Ltd./sanofi-Aventis.

A. Rovira serves on scientific advisory boards for Biogen Idec, Novartis, Sanofi-Genzyme, and OLEA Medical, has received speaker honoraria from Bayer, Sanofi-Genzyme, Bracco, Merck-Serono, Teva Pharmaceutical Industries Ltd, Novartis and Biogen Idec, and has research agreements with Siemens AG and Icometrix.

M.A. Rocca received speaker’s honoraria from Biogen Idec, Novartis, TEVA, Genzyme and ExceMed and receives research support from the Italian Ministry of Health and Fondazione Italiana Sclerosi Multipla.

M. Filippi is Editor-in-Chief of the Journal of Neurology; serves on scientific advisory board for Teva Pharmaceutical Industries; has received compensation for consulting services and/or speaking activities from Biogen Idec, Excemed, Novartis, and Teva Pharmaceutical Industries; and receives research support from Biogen Idec, Teva Pharmaceutical Industries, Novartis, Italian Ministry of Health, Fondazione Italiana Sclerosi Multipla, Cure PSP, Alzheimer’s Drug Discovery Foundation (ADDF), the Jacques and Gloria Gossweiler Foundation (Switzerland), and ARiSLA (Fondazione Italiana di Ricerca per la SLA).

H. Vrenken has received research grants from Pfizer, MerckSerono, Novartis and Teva, and speaker honoraria from Novartis and MerckSerono; all funds were paid directly to his institution.

CAM Gandini Wheeler-Kingshott receives research grants (PI and co-applicant) from ISRT, EPSRC, Wings for Life, UK MS Society, Horizon2020, Biogen and Novartis.

L Pirpamer has nothing to disclose.

Declan Chard has received honoraria (paid to his employer) from Ismar Healthcare NV, Swiss MS Society, Excemed (previously Serono Symposia International Foundation), Merck, Bayer and Teva for faculty-led education work; Teva for advisory board work; meeting expenses from Merck, Teva, Novartis, the MS Trust and National MS Society; and has previously held stock in GlaxoSmithKline.

AJ Thompson receives grant support from the Multiple Sclerosis Society of Great Britain and Northern Ireland, and has received honoraria/support for travel for consultancy from Eisai, Biogen (Optum Insight), and Excemed. He received support for travel from the International Progressive MS Alliance, as chair of their Scientific Steering Committee and the National MS Society (USA) as member of their Research Programs Advisory Committee, and receives an honorarium from SAGE Publishers as Editor-in-Chief of Multiple Sclerosis Journal.

O Ciccarelli receives research grant support from the Multiple Sclerosis Society of Great Britain and Northern Ireland, the NIHR UCLH Biomedical Research Centre; she is a consultant for Teva, Roche, Novartis, Biogen, Genzyme and GE. She is an Associate Editor for Neurology, for which she receives an honorarium.

AE, FP, DCA, WB, DRA, CT, MJC, FDA, SHP, NC, MLS, MB, SR, CG, JSG, CEL, LP, CE, SO, have nothing to report in relation to this study.

## Abbreviations

SD: standard deviation;
CIS: clinically isolated syndrome;
RRMS: relapsing-remitting multiple sclerosis;
SPMS: secondary-progressive multiple sclerosis;
PPMS: primary-progressive multiple sclerosis;
ml: millilitre;
EDSS: expanded-disability status scale;
DMTs: disease modifying treatment.

## References

1. Browne P, Chandraratna D, Angood C, et al Atlas of Multiple Sclerosis 2013: A growing global problem with widespread inequity. Neurology 2014;83(11):1022–1024.

2. Mahad DH, Trapp BD, Lassmann H. Pathological mechanisms in progressive multiple sclerosis. Lancet Neurol. 2015;14(2):183–193.

3. Geurts JJG, Calabrese M, Fisher E, Rudick RA. Measurement and clinical effect of grey matter pathology in multiple sclerosis. Lancet Neurol. 2012;11(12):1082–1092.

4. Bermel RA, Bakshi R. The measurement and clinical relevance of brain atrophy in multiple sclerosis. Lancet Neurol. 2006;5(2):158–170.

5. De Stefano N, Giorgio A, Battaglini M, et al Assessing brain atrophy rates in a large population of untreated multiple sclerosis subtypes. Neurology 2010;74(23):1868–1876.

6. De Stefano N, Stromillo ML, Giorgio A, et al Establishing pathological cut-offs of brain atrophy rates in multiple sclerosis. J. Neurol. Neurosurg. Psychiatry 2015;jnnp-2014-309903.

7. Chataway J, Schuerer N, Alsanousi A, et al Effect of high-dose simvastatin on brain atrophy and disability in secondary progressive multiple sclerosis (MS-STAT): a randomised, placebo-controlled, phase 2 trial. Lancet Lond. Engl. 2014;383(9936):2213–2221.

8. Chataway J. MS-SMART: Multiple Sclerosis-Secondary Progressive Multi-Arm Randomisation Trial - ClinicalTrials.gov [Internet]. [date unknown];[cited 2016 Oct 20] Available from: https://clinicaltrials.gov/ct2/show/study/NCT01910259?term=mssmart&rank=1

9. Fisher E, Lee J-C, Nakamura K, Rudick RA. Gray matter atrophy in multiple sclerosis: a longitudinal study. Ann. Neurol. 2008;64(3):255–265.

10. Fisniku LK, Brex PA, Altmann DR, et al Disability and T2 MRI lesions: a 20-year follow-up of patients with relapse onset of multiple sclerosis. Brain J. Neurol. 2008;131(Pt 3):808–817.

11. Roosendaal SD, Bendfeldt K, Vrenken H, et al Grey matter volume in a large cohort of MS patients: relation to MRI parameters and disability. Mult. Scler. J. 2011;17(9):1098–1106.

12. Steenwijk MD, Geurts JJG, Daams M, et al Cortical atrophy patterns in multiple sclerosis are non-random and clinically relevant. Brain J. Neurol. 2016;139(Pt 1):115–126.

13. Eshaghi A, Bodini B, Ridgway GR, et al Temporal and spatial evolution of grey matter atrophy in primary progressive multiple sclerosis. NeuroImage 2014;86:257–264.

14. Rocca MA, Mesaros S, Pagani E, et al Thalamic damage and long-term progression of disability in multiple sclerosis. Radiology 2010;257(2):463–469.

15. Schoonheim MM, Hulst HE, Brandt RB, et al Thalamus structure and function determine severity of cognitive impairment in multiple sclerosis. Neurology 2015;84(8):776–783.

16. Henneman WJP, Sluimer JD, Barnes J, et al Hippocampal atrophy rates in Alzheimer disease: added value over whole brain volume measures. Neurology 2009;72(11):999–1007.

17. Ceccarelli A, Rocca MA, Pagani E, et al A voxel-based morphometry study of grey matter loss in MS patients with different clinical phenotypes. NeuroImage 2008;42(1):315–322.

18. Polman CH, Reingold SC, Banwell B, et al Diagnostic criteria for multiple sclerosis: 2010 revisions to the McDonald criteria. Ann. Neurol. 2011;69(2):292–302.

19. Lublin FD, Reingold SC, Cohen JA, et al Defining the clinical course of multiple sclerosis: the 2013 revisions. Neurology 2014;83(3):278–286.

20. Kurtzke JF. Rating neurologic impairment in multiple sclerosis: an expanded disability status scale (EDSS). Neurology 1983;33(11):1444–1452.

21. Tustison NJ, Avants BB, Cook PA, et al N4ITK: improved N3 bias correction. IEEE Trans. Med. Imaging 2010;29(6):1310–1320.

22. Schmidt P, Gaser C, Arsic M, et al An automated tool for detection of FLAIR-hyperintense white-matter lesions in Multiple Sclerosis. NeuroImage 2012;59(4):3774–3783.

23. Jenkinson M, Smith S. A global optimisation method for robust affine registration of brain images. Med. Image Anal. 2001;5(2):143–156.

24. Zhang Y, Brady M, Smith S. Segmentation of brain MR images through a hidden Markov random field model and the expectation-maximization algorithm. IEEE Trans Med Imaging 2001;20(1):45–57.

25. Battaglini M, Jenkinson M, De Stefano N. Evaluating and reducing the impact of white matter lesions on brain volume measurements. Hum. Brain Mapp. 2012;33(9):2062–2071.

26. Popescu V, Agosta F, Hulst HE, et al Brain atrophy and lesion load predict long term disability in multiple sclerosis. J. Neurol. Neurosurg. Psychiatry 2013;84(10):1082–1091.

27. Amato MP, Hakiki B, Goretti B, et al Association of MRI metrics and cognitive impairment in radiologically isolated syndromes. Neurology 2012;78(5):309–314.

28. Reuter M, Fischl B. Avoiding asymmetry-induced bias in longitudinal image processing. NeuroImage 2011;57(1):19–21.

29. Reuter M, Rosas HD, Fischl B. Highly accurate inverse consistent registration: a robust approach. NeuroImage 2010;53(4):1181–1196.

30. Reuter M, Schmansky NJ, Rosas HD, Fischl B. Within-subject template estimation for unbiased longitudinal image analysis. NeuroImage 2012;61(4):1402–1418.

31. Cardoso MJ, Modat M, Wolz R, et al Geodesic information flows: spatially-variant graphs and their application to segmentation and fusion. IEEE Trans. Med. Imaging 2015;34(9):1976–1988.

32. Klein A, Tourville J. 101 labeled brain images and a consistent human cortical labeling protocol [Internet]. Front. Neurosci. 2012;6[cited 2016 Jul 7] Available from: http://journal.frontiersin.org/article/10.3389/fnins.2012.00171/abstract

33. Pardini M, Sudre CH, Prados F, et al Relationship of grey and white matter abnormalities with distance from the surface of the brain in multiple sclerosis. J. Neurol. Neurosurg. Psychiatry 2016;87(11):1212–1217.

34. Bocchetta M, Cardoso MJ, Cash DM, et al Patterns of regional cerebellar atrophy in genetic frontotemporal dementia. NeuroImage Clin. 2016;11:287–290.

35. Prados F, Cardoso MJ, Leung KK, et al Measuring brain atrophy with a generalized formulation of the boundary shift integral. Neurobiol. Aging 2015;36 Suppl 1:S81–90.

36. Jorge Cardoso M, Leung K, Modat M, et al STEPS: Similarity and Truth Estimation for Propagated Segmentations and its application to hippocampal segmentation and brain parcelation. Med. Image Anal. 2013;17(6):671–684.

37. Bernal-Rusiel JL, Greve DN, Reuter M, et al Statistical analysis of longitudinal neuroimage data with Linear Mixed Effects models. NeuroImage 2013;66:249–260.

38. Vittinghoff E, Glidden DV, Shiboski SC, McCulloch CE. Regression Methods in Biostatistics: Linear, Logistic, Survival, and Repeated Measures Models [Internet]. Springer New York; 2006.Available from: https://books.google.co.uk/books?id=tGw-9HRV2UEC

39. Zhang D, Raichle ME. Disease and the brain’s dark energy. Nat. Rev. Neurol. 2010;6(1):15–28.

40. R Core Team. R: A Language and Environment for Statistical Computing [Internet]. Vienna, Austria: R Foundation for Statistical Computing; 2014.Available from: http://www.R-project.org/

41. Pinheiro JC, Bates D. Mixed-Effects Models in S and S-PLUS [Internet]. Springer; 2009. Available from: https://books.google.co.uk/books?id=y54QDUTmvDcC

42. Healy BC, Engler D, Glanz B, et al Assessment of definitions of sustained disease progression in relapsing-remitting multiple sclerosis. Mult. Scler. Int. 2013;2013:189624.

43. Sabuncu MR, Bernal-Rusiel JL, Reuter M, et al Event time analysis of longitudinal neuroimage data. NeuroImage 2014;97:9–18.

44. Therneau TM, Grambsch PM. Modeling Survival Data: Extending the Cox Model [Internet]. Springer New York; 2013. Available from: https://books.google.co.uk/books?id=oj0mBQAAQBAJ

45. Zivadinov R, Heininen-Brown M, Schirda CV, et al Abnormal subcortical deep-gray matter susceptibility-weighted imaging filtered phase measurements in patients with multiple sclerosis: a case-control study. NeuroImage 2012;59(1):331–339.

46. Kolasinski J, Stagg CJ, Chance SA, et al A combined post-mortem magnetic resonance imaging and quantitative histological study of multiple sclerosis pathology. Brain J. Neurol. 2012;135(Pt 10):2938–2951.

47. Hametner S, Wimmer I, Haider L, et al Iron and neurodegeneration in the multiple sclerosis brain. Ann. Neurol. 2013;74(6):848–861.

48. Bodini B, Chard D, Altmann DR, et al White and gray matter damage in primary progressive MS: The chicken or the egg? Neurology 2016;86(2):170–176.

49. Howell OW, Reeves CA, Nicholas R, et al Meningeal inflammation is widespread and linked to cortical pathology in multiple sclerosis. Brain J. Neurol. 2011;134(Pt 9):2755–2771.

50. Haider L, Zrzavy T, Hametner S, et al The topograpy of demyelination and neurodegeneration in the multiple sclerosis brain. Brain J. Neurol. 2016;139(Pt 3):807–815.

51. Lindberg RLP, De Groot CJA, Certa U, et al Multiple sclerosis as a generalized CNS disease-comparative microarray analysis of normal appearing white matter and lesions in secondary progressive MS. J. Neuroimmunol. 2004;152(1–2):154–167.

52. Serafini B, Rosicarelli B, Magliozzi R, et al Detection of ectopic B-cell follicles with germinal centers in the meninges of patients with secondary progressive multiple sclerosis. Brain Pathol. Zurich Switz. 2004;14(2):164–174.

53. Seeley WW, Crawford RK, Zhou J, et al Neurodegenerative diseases target large-scale human brain networks. Neuron 2009;62(1):42–52.

54. Pellicano C, Gallo A, Li X, et al Relationship of cortical atrophy to fatigue in patients with multiple sclerosis. Arch. Neurol. 2010;67(4):447–453.

55. Kutzelnigg A, Lucchinetti CF, Stadelmann C, et al Cortical demyelination and diffuse white matter injury in multiple sclerosis. Brain J. Neurol. 2005;128(Pt 11):2705–2712.

56. Audoin B, Zaaraoui W, Reuter F, et al Atrophy mainly affects the limbic system and the deep grey matter at the first stage of multiple sclerosis. J. Neurol. Neurosurg. Psychiatry 2010;81(6):690–695.

57. Kalkers NF, Ameziane N, Bot JC, et al Longitudinal brain volume measurement in multiple sclerosis: rate of brain atrophy is independent of the disease subtype. Arch. Neurol. 2002;59(10):1572–1576.

58. Lukas C, Minneboo A, de Groot V, et al Early central atrophy rate predicts 5 year clinical outcome in multiple sclerosis. J. Neurol. Neurosurg. Psychiatry 2010;81(12):1351–1356.

59. Riccitelli G, Rocca MA, Pagani E, et al Cognitive impairment in multiple sclerosis is associated to different patterns of gray matter atrophy according to clinical phenotype. Hum. Brain Mapp. 2011;32(10):1535–1543.

60. Mallik S, Muhlert N, Samson RS, et al Regional patterns of grey matter atrophy and magnetisation transfer ratio abnormalities in multiple sclerosis clinical subgroups: a voxel-based analysis study. Mult. Scler. Houndmills Basingstoke Engl. 2015;21(4):423–432.

61. Lansley J, Mataix-Cols D, Grau M, et al Localized grey matter atrophy in multiple sclerosis: a meta-analysis of voxel-based morphometry studies and associations with functional disability. Neurosci. Biobehav. Rev. 2013;37(5):819–830.

62. Filippi M, Rovaris M, Inglese M, et al Interferon beta-1a for brain tissue loss in patients at presentation with syndromes suggestive of multiple sclerosis: a randomised, double-blind, placebo-controlled trial. Lancet Lond. Engl. 2004;364(9444):1489–1496.

63. Kappos L, Edan G, Freedman MS, et al The 11-year long-term follow-up study from the randomized BENEFIT CIS trial. Neurology 2016;

64. Biberacher V, Schmidt P, Keshavan A, et al Intra- and interscanner variability of magnetic resonance imaging based volumetry in multiple sclerosis. NeuroImage 2016;

65. Bendfeldt K, Hofstetter L, Kuster P, et al Longitudinal gray matter changes in multiple sclerosis-differential scanner and overall disease-related effects. Hum. Brain Mapp. 2012;33(5):1225–1245.

66. Smith SM, Zhang Y, Jenkinson M, et al Accurate, robust, and automated longitudinal and cross-sectional brain change analysis. NeuroImage 2002;17(1):479–489.

67. Smith SM, Rao A, De Stefano N, et al Longitudinal and cross-sectional analysis of atrophy in Alzheimer’s disease: cross-validation of BSI, SIENA and SIENAX. NeuroImage 2007;36(4):1200–1206.

68. Cash DM, Frost C, Iheme LO, et al Assessing atrophy measurement techniques in dementia: Results from the MIRIAD atrophy challenge. NeuroImage 2015;123:149–164.

69. Frost C, Kenward MG, Fox NC. The analysis of repeated “direct“ measures of change illustrated with an application in longitudinal imaging. Stat. Med. 2004;23(21):3275–3286.

70. Malone IB, Leung KK, Clegg S, et al Accurate automatic estimation of total intracranial volume: a nuisance variable with less nuisance. NeuroImage 2015;104:366–372.

71. Hedman AM, van Haren NE, Schnack HG, et al Human brain changes across the life span: a review of 56 longitudinal magnetic resonance imaging studies. Hum Brain Mapp 2012;33(8):1987–2002.

72. Fraser MA, Shaw ME, Cherbuin N. A systematic review and meta-analysis of longitudinal hippocampal atrophy in healthy human ageing. NeuroImage 2015;112:364–374.

73. University of California, San Francisco MS-EPIC Team:, Cree BAC, Gourraud P-A, et al. Long-term evolution of multiple sclerosis disability in the treatment era. Ann. Neurol. 2016;

